# NetVA: An R Package for Network Vulnerability and Influence Analysis

**DOI:** 10.1101/2023.07.31.551200

**Authors:** Swapnil Kumar, Grace Pauline, Vaibhav Vindal

## Abstract

In biological network analysis, identifying key molecules plays a decisive role in the development of potential diagnostic and therapeutic candidates. Among various approaches of network analysis, network vulnerability analysis is quite important, as it assesses significant associations between topological properties and the functional essentiality of a network. Further, some node centralities are also used to screen out key molecules. Among these node centralities, escape velocity centrality (EVC), and its extended version (EVC+) outperform others, *viz*., Degree, Betweenness, and Clustering coefficient. Keeping this in mind, we aimed to develop a first-of-its-kind R package named NetVA, which analyzes networks to identify key molecular players through network vulnerability and EVC+-based approaches. To demonstrate the application and relevance of our package in network analysis, previously published and publicly available protein-protein interactions (PPIs) data of human breast cancer were analyzed. This resulted in identifying some most important proteins. These included essential proteins, non-essential proteins, hubs, and bottlenecks, which play vital roles in breast cancer development. Thus, the NetVA package, available at https://github.com/kr-swapnil/NetVA with a detailed tutorial to download and use, assists in predicting potential candidates for therapeutic and diagnostic purposes by exploring various topological features of a disease-specific PPIs network.

## Introduction

The genome-wide interactions of genes or gene products lead to the formation of complex systems like structures known as biological networks. Because biological molecules such as proteins are known to perform collectively for their functions and metabolic pathways by interacting with each other.^1,2^ These networks can be disease-, process- or species-specific. Any alteration in a network system of a normal state, e.g., deletion, insertion, or inactivation of nodes, i.e., proteins, results in a disrupted or collapsed system with altered normal processes and functions, which in turn leads to the development of a diseased state.^3-5^ For example, in the case of bacterial or viral disease, the pathogen’s proteins interact with the host’s proteins and alter the host’s protein-protein interactions network (PPIN). The prediction of key molecules that can lead to the reduced robustness or increased vulnerability of systems with capabilities of transforming healthy cells from normal conditions to disease conditions is of great importance; because it helps to estimate and understand the effect of targeted perturbations such as drug-mediated inactivation on underlying physiological and pathological states of diseased systems.

The development of computational models representing complex biological systems of interest and subsequent analysis has been utilized previously for a thorough exploration and understanding of underlying mechanisms and related properties.^6-9^ The analysis and exploration of these complex networks lead to identifying key molecular players having decisive roles in cellular and functional activities. These key molecules, e.g., genes or proteins, may aid in the development of potential diagnostic and therapeutic targets.

Among various approaches of network analysis, network vulnerability analysis is quite important, as evident from several previous works.^10-13^ Network vulnerability analysis is a method of estimating structural changes leading to substantial changes in functions caused by dysregulated or altered components of the network. In general, the dysregulation or alteration of network components or networks can be achieved either by deleting nodes or rewiring their connections. In other words, it encompasses assessments of significant associations between topological properties and the functional essentiality of the network. There is a publicly available tool named eNelator^14^ for this purpose. However, the eNelator, freely available from the authors on request (currently unavailable), is a Java-based Cytoscape plugin that does not support parallel processing. Thus, analyzing networks using this plugin is time-consuming, especially for networks of large size. Further, only a few topological properties are available in this Cytoscape plugin for analysis. Table 1 provides an overview of the functionality and different features these software/packages support.

**Table 1.** Comparison of NetVA with eNelator Software.

| Software/Package | Type | Topological features | Escape velocity centrality<sup>a</sup> | Parallel computing | Working status |
| --- | --- | --- | --- | --- | --- |
| NetVA | R package | 11 | Yes | Yes | Fully functional |
| eNelator | Cytoscape app (Java-based) | 5 | No | No | Not known |

<sup>a</sup> Not available in any software or packages till now except NetVA.

Besides various methods of network analysis based on local and global topological features, several other methods have also been proposed to identify key nodes in complex networks. Out of these approaches, the escape velocity centrality (EVC) and its extended version (EVC+), recently proposed node centrality measures based on escape velocity formulae, provide the most effective ranking method to identify the most influential nodes (INs) based on their positions and connections in the network.^15^

Keeping the importance of network vulnerability analysis, the effectiveness of EVC, and the limitations of related tools in mind, we aim to develop an R package named NetVA, which performs network analysis using two approaches: (i) network vulnerability and (ii) EVC to identify key nodes. To demonstrate the application and effectiveness of our package in network analysis, previously published data on protein-protein interactions (PPIs) of human breast cancer were analyzed. Consequently, it revealed some most vulnerable proteins (VPs) and influential proteins (IPs) consistent with previously reported experimental evidence, out of which few were either essential, hub, or bottleneck proteins. At the same time, others were neither essential, hub, nor bottleneck proteins. Hub proteins in cellular networks are most important for the survival of cells and hence, are essential proteins.^16^ In other words, a gene or protein is essential when the loss of its function leads to the compromised fitness or viability of the organism.^17^ Thus, the package can assist in identifying putative diagnostic and therapeutic candidates by exploring various network properties through vulnerability analysis at the systems level.

## Methods

### Protein-protein interactions data

The protein-protein interactions (PPIs) data of breast cancer were downloaded for the network analysis from the CancerNet database.^18^ The CancerNet is a database of cancer-associated information on PPIs, functionally synergistic pairs of miRNA–miRNA and miRNA–target interactions across 33 types of human cancers. Additionally, information about PPIs across 33 main types of normal tissues and cells is also included in the database.

### Breast cancer-associated and essential genes

A comprehensive list of breast cancer-associated genes was obtained by combining two lists of breast cancer genes, one from a previous network-based study on breast cancer^7^ and another from a data curation study on breast cancer genes^19,20^ compiled from different sources. A list of essential genes was also retrieved from previously published literature.^17^ These lists of breast cancer-associated and essential genes were used for the subsequent analysis.

### Implementation and development

The methods of network vulnerability and influence analysis were implemented using R (www.r-project.org)^21^ and developed a software package named ‘NetVA’ for this purpose. R is an open-source software environment implementing various base functions and also a programming language for statistical computing. It provides the most sophisticated computing platform for the exploration, analysis, and annotation of multiple types of biological data, including gene expression profiles, DNA/protein sequences, and biological networks, with the help of various R/Bioconductor packages such as limma,^22^ DESeq2,^23^ edgeR,^24^ seqinR,^25^ and igraph.^26^ Besides the base functions of R, the igraph R package was also used in developing the NetVA package.

Further, to demonstrate the application of the NetVA package in network analysis, PPIs data of breast cancer were used to reconstruct a network. Subsequently, the network was analyzed by deleting all nodes present in the network one by one and constructing the resultant network each time. This followed the calculation of various topological properties. Thus, the analysis resulted in lists of values of the calculated properties for each of the deleted nodes, which provided screening criteria for the VPs of the network; thus, the analysis led to the identification of a list of key molecules based on the criteria of more than two topological properties. These VPs were also checked for their presence in the list of essential genes.

Additionally, the network was analyzed by deleting all node pairs (edges) and node triplets (three nodes connected together at least with two edges) present in the network one by one and reconstructing the resultant network each time. This also followed the calculation of various topological properties of the networks. In both conditions (node pairs and node triplets), the analysis resulted in lists of values of the calculated properties for each of the deleted node pairs and node triplets. Thus, the analysis reveals the network’s most critical or vulnerable protein pairs (VPPs) and vulnerable protein triplets (VPTs) based on these properties (more than two) as screening criteria.

The network was also analyzed using EVC and its extended version (EVC+), as proposed by Ullah et al. in their recent study.^15^ The EVC and EVC+ provided new centrality-based criteria to screen out key nodes by ranking them based on their position (K-shell and shortest paths) in the network and the number of connections (degree) they have with their neighboring nodes. The EVC and EVC+-based ranking of nodes has been demonstrated to outperform other node centralities such as Degree, Betweenness, and Clustering coefficient in identifying key INs. Further, the average values of EVC and EVC+ of the top 20% of nodes were considered as cutoff values to identify IPs using the Pareto principle of the 80:20 rule.^27^ Subsequently, these two lists of IPs were cross-checked with the list of protein products of essential genes.

### Topological properties of a network

A network’s topological properties measure its overall robustness and help to understand its structural, functional, and behavioral organization. The topological properties are helpful for finding critical or vulnerable nodes (VNs), node pairs (VNPs), and node triplets (VNTs) in the network. The VNs, VNPs, and VNTs are identified by knocking down each node, node pair (edge), and node triplet from the network, respectively, one by one and calculating the topological properties of the resultant network. Thus, the topological properties play a significant role in identifying potential drug targets for various diseases. The topological or network properties that can be calculated for a given network using the NetVA include average node connectivity, average closeness, articulation point, average path length, average betweenness centrality, clustering coefficient, diameter, global efficiency, network centralization, network density, and network heterogeneity. The calculated values of these properties for each node, node pair, and node triplet can be used to identify VPs among all proteins considered for the analysis.

A node, node pair, or node triplet can be considered important if the deletion or removal of that node, node pair, or node triplet from the network leads to the high values of articulation point, average path length, and diameter of that network, whereas the low values of average node connectivity, average betweenness centrality, clustering coefficient, average closeness centrality, global efficiency, network centralization, network density, and heterogeneity. All the network properties studied in the present work have been discussed further in detail as defined previously.^28^

#### Average node connectivity

The average node connectivity or degree (ANC) is the mean value of degree values of all nodes in a network of interest. Highly connected genes, i.e., hub genes that have major contributions to the value of ANC, play vital roles in the behavioral organization of biological networks.^16,29-31^ Additionally, degree or connectivity has been found to be an important parameter for screening and identifying biologically significant genes in diseases, including cancer.^6,32^

#### Articulation point

An articulation point (AP) is a node whose removal from a network causes the network to split into different sub-networks. Thus, the APs cause alteration in the network connectivity and act as VNs of the network because their deletion can increase the count of disconnected components or sub-networks.^33^ These APs can be identified by implementing a brute force approach where a network’s connectivity is checked after each node’s removal.

#### Average path length

The average path length (APL) determines the overall connectivity of the network. In other words, it is the value of the average shortest path among all pairs of nodes present in the network.

#### Betweenness centrality

The betweenness centrality (BC) is defined as the property of a node being a connector of two or more clusters in a given network. The BC is the number of all shortest paths that pass through the node of interest in a given network.

#### Closeness centrality

The closeness centrality (CC) of a node is denoted as the reciprocal of the sum of distances between that node to all the other nodes in a network.^34^ It measures the number of steps required to access all other nodes from a given node.

#### Clustering coefficient

The clustering coefficient (CC) can be defined as the ratio of the total number of edges a node has and the number of all possible edges a node can have. It determines the local interconnectivity of the network as it measures whether neighbors of a particular node are neighbors of each other. The CC of a node is the density measure of cliquishness or local connections.^35,36^

#### Diameter

The diameter or network diameter (NDR) is the shortest path length between the two most distantly placed nodes in a given network. In other words, the diameter is the longest of all the shortest path lengths calculated from every node to all other nodes in the network.

#### Global efficiency

A network’s global efficiency (GE) is the average of reciprocal distances between all pairs of nodes in the network.

#### Network centralization

The network centralization (NC) or simply centralization (also known as degree centralization) of a network is a basic index used widely for connectivity distribution.^34^ It provides an idea about specific nodes being far more central than others within the network of interest. The centralization index can be used to describe differences between networks at the structural level, e.g., structural differences among metabolic networks.^37^

#### Network density

The network density (ND) can be defined as the ratio of the number of observed edges to the number of all possible edges in a given network. In other words, it is the actual number of connections divided by the possible number of connections. A completely connected network will have a density of one. In contrast, others will have less than one, i.e., a decimal value representing the percentage value of possible connections actually present in the network.

#### Network heterogeneity

The network heterogeneity (NH) represents the tendency of a network to have a very heterogeneous nature, i.e., some highly connected (hub) nodes and the majority of nodes with very few connections. It is the ratio of the standard deviation of the degree to the average degree value.

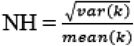

Where k is the node degree in a network.

### Identification of hub and bottleneck proteins

From the PPIN of breast cancer, we identified a list of hubs and bottlenecks to check whether these identified VPs are hubs and bottlenecks. For this, the Pareto principle of the 80:20 rule^27^ was followed. The average value of degrees and the average value of betweenness of the top 20% of nodes were considered cutoff values to identify hubs and bottlenecks, respectively. Further, these lists of hubs and bottlenecks were cross-checked with the list of VPs to identify VPs with hub and bottleneck properties.

## Results

### Network vulnerability assessment

The vulnerability of a network can be assessed by calculating and comparing its APL with the APL of the resultant or mutated networks constructed after deleting one or more nodes. Herein, we first deleted all nodes from the network one by one using single-node and two-node deletion approaches independently. Subsequently, the effect of deletion of one or more than one node on the stability of the original structure of the network was measured globally. Thus, it provided an approximation of how vulnerable the given network is to the removal or disruption of specific proteins or groups of proteins. A protein can be defined as vulnerable if after removal of that from the network under investigation causes major changes in the APL of that network. When we investigated the APL of original and mutated networks after removing proteins using both single and two-node approaches, it was found that there were significant changes in their relative values (**Table 2**), which signifies the importance of network vulnerability analysis to identify key VPs.

**Table 2.** Distribution of APL before and after removal of nodes from the network.

| Approach | WN-APL | MN-APL | RAPL |
| --- | --- | --- | --- |
| Single Node Deletion | 2.380426 | 2.38048 | 1.000023 |
| Two Node Deletion | 2.380426 | 2.382536 | 1.000886 |

WN-APL, APL of the original network; MN-APL, APL (average value) of all the resultant/mutated network after removal of nodes one by one; RAPL, relative APL of the mutated network relative to the original network.

### Software/package description

Herein, we describe the structure, functionalities, and features of the NetVA package. The software package implemented in R is available at https://github.com/kr-swapnil/NetVA with a detailed tutorial. The NetVA package depends on the base functions of R, parallel R package integrated with base R, and igraph R package. It runs under major operating systems like Microsoft Windows, macOS, and Linux.

The NetVA package contains a set of functions for network analysis, including the simulation and construction of a PPIs network after deleting one node (protein), node pair, or node triplet at a time one by one from the original network for (i) n times, if the total number of nodes to be deleted in the network, is n, where n is the total number of nodes in the network; or (ii) m times if the user wants to check the effect of deletion of only selected nodes, i.e., m nodes on the given network. Functions in the NetVA package can be divided into four main categories: (i) Calculation of global properties of a given network after deletion of each node, node pair, or node triplet at a time one by one as specified by the user; (ii) Exporting the results of properties calculation for further use; (iii) Detection of VPs, VPPs, or VPTs based on the five-point summary of the boxplot; and (iv) Calculation of values of EVC and EVC+ for all nodes (proteins) and identifying INs based on these values. All these functions of the package have been described below, and a functional overview has been depicted in Figure 1.

1. netva: it is a function that performs network vulnerability analysis in normal mode on Windows-based machines and parallel mode using multiple cores on Linux/macOS machines. It deletes one or more nodes (e.g., two nodes, i.e., node pair, three nodes, i.e., node triplet) at once and constructs the resultant network having (n-m) nodes, where n is the total number of nodes and m is the number of nodes deleted from the original network. This is followed by the calculation of 11 different topological properties as described in the Methods section. This step is repeated for all elements of the input vector or list of nodes, node pairs, or node triplets.
2. evc: it is a function to calculate EVC and EVC+ values of all nodes in a given network.
3. heterogeneity: it calculates the value of network heterogeneity for a given network as input by the user.
4. detectVNs: it detects VNs, VNPs, or VNTs based on a five-point summary of the boxplot and lower or higher values as discussed in the Methods section of each calculated topological property for the resultant network.
5. detectHubs: it detects hub nodes based on the Pareto principle of 80-20 by using the average degree value of the top 20% of nodes as the cutoff.
6. detectBottlenecks: it detects bottleneck nodes based on the Pareto principle of 80-20 by using the average betweenness value of the top 20% of nodes as the cutoff.
7. detectINs: it detects INs based on the Pareto principle of 80-20 by using the average values of EVC and EVC+ of the top 20% of nodes as the cutoff.

**Figure 1.**
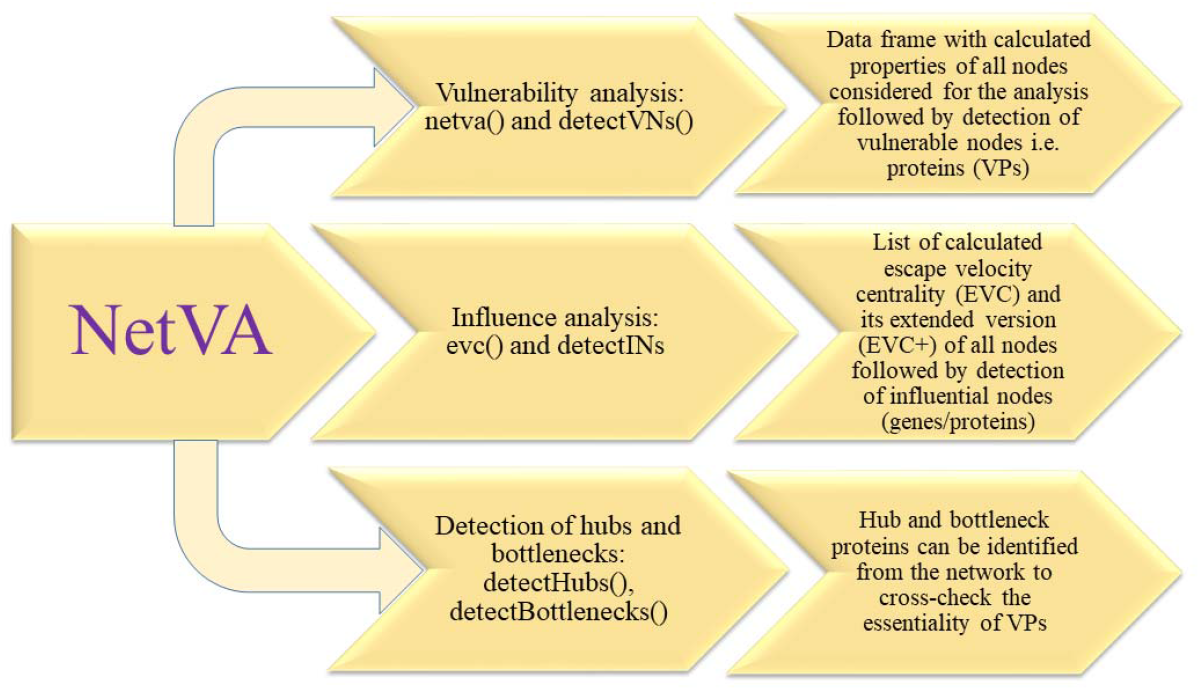
Functional overview of the NetVA R package.

The NetVA package simulates an attack on PPIN by both ways measuring the global effects on the stability of PPIN in terms of several topological measurements of the network as discussed in the Methods section, including average path length, articulation point, average node connectivity, betweenness centrality, clustering coefficient, network centralization, density, diameter, and heterogeneity. The second application of this package is to identify IPs based on EVC and EVC+ values of nodes in the given network.

A network may be connected entirely or may not be. If it is not fully connected, it can have one large connected component and a few small to medium-sized connected components. In this case, the user may select only the large connected component of the network while performing the analysis to identify VPs, protein pairs, or protein triplets present therein.

### Application: breast cancer network

We applied NetVA on publicly available PPIs data of breast cancer from the CancerNet database as a case study to demonstrate the application of NetVA for network vulnerability and influence analysis. The PPIs data of breast cancer used in this study consisted of 143751 interactions (edges) among 10597 proteins (nodes) which was used to construct a breast cancer network for subsequent use in the present analysis.

#### Vulnerable proteins (VPs)

All 10597 nodes in the breast cancer network were considered for the analysis using the NetVA package. One by one, these nodes were deleted from the original network, and the resultant networks were constructed. It followed the calculation of different topological features of these networks (**Table S1**). Further, all VPs present in the network were screened out based on the 11 calculated topological features (**Table S1**). There were 1107 unique VPs based on a minimum of three properties (details given in **Table S2**). Subsequently, the analysis of breast cancer PPIN for identifying hubs and bottlenecks revealed 519 hub and 156 bottleneck proteins. Among all VPs, it was noticed that there were only two VPs, i.e., SUMO2 and UBC, as per eight properties, and these were also hubs and bottlenecks (**Table S2**).

#### Influence analysis and influential proteins (IPs)

As discussed in the Methods section, the EVC and EVC+-based influence analysis of breast cancer PPIN revealed 704 and 701 IPs, respectively. Thus, 656 unique IPs were commonly present in the EVC and EVC+-based lists of IPs (**Table S3**). Of these common IPs, 91 proteins were products of essential genes. When we intersected the list of 1107 VPs with the list of 656 IPs, 646 unique proteins were found as vulnerable cum influential proteins (VIPs) (**Table S4**).

As discussed in the Methods section, these VPs, IPs, and VIPs were further checked for their essentiality in terms of hub-ness (degree) and connected-ness (betweenness). These VPs were also matched with protein products of 1103 essential genes^17^ to check their essentiality. Further, the list of breast cancer genes (obtained by combining two lists, one with 617 genes and the other with 956 genes) consisted of 1260 unique genes. These breast cancer genes were also used to check the importance and involvement of VPs in breast cancer.

Out of 1107 VPs, there were 516 hubs, 156 bottlenecks, 199 proteins as products of breast cancer genes, and 143 proteins as products of essential genes (**Table 3**). Further, among 656 IPs, there were 514 hubs, 147 bottlenecks, 131 proteins as products of breast cancer genes, and 91 proteins as products of essential genes. Similarly, among 646 VIPs, there were 511 hubs, 147 bottlenecks, 131 proteins as products of breast cancer genes, and 91 proteins as products of essential genes.

**Table 3.** Distribution of vulnerable and influential proteins.

| **Protein** | **Total** | **Hub** | **Bottleneck** | **BCG** | **EG** |
| --- | --- | --- | --- | --- | --- |
| VP | 1107 | 516 | 156 | 199 | 143 |
| IP | 656 | 514 | 147 | 131 | 91 |
| VIP | 646 | 511 | 147 | 131 | 91 |

VP: Vulnerable proteins; IP: Influential proteins; VIP: Vulnerable cum influential proteins; BCG: Breast cancer genes; EG: Essential genes.

#### Vulnerable protein pairs (VPPs)

A total of 4749 protein pairs (edges) involving 646 unique VIPs in the network were analyzed by deleting them individually and reconstructing the resultant network using two node deletion approach to identify VPPs. It calculated 11 different topological properties of the 4749 resultant networks (**Table S5**). Thus, it resulted in 582 VPPs based on at least three calculated properties of the resultant network (**Table S6**). Out of these, 16 VPPs *viz*. ACTN1-UBC, EIF3A-UBC, ESR1-UBC, HDGF-UBC, HSPA1A-UBC, HSPB1-UBC, HUWE1-UBC, IKBKB-UBC, LYN-UBC, MAPK6-UBC, MET-UBC, PAN2-UBC, PCBP1-UBC, RBM14-UBC, SRPK2-UBC, and UBB-UBC were most vulnerable based on a maximum of eight properties. Further, 14 VPPs out of 582 viz. ATM-MDM2, BCL2-MYC, CDK4-MDM2, CDK4-TP53, ETS1-TP53, MET-TP53, MTOR-MYC, MTOR-TP53, PLK1-TP53, RAD21-TP53, RELA-TP53, RPS12-TP53, SKIL-TP53, and TOP1-TP53 were synthetic lethal pairs.

## Discussion

The vulnerability analysis is a method of analyzing networks that estimates structural changes in a network leading to substantial changes in its functions caused by any dysregulation or alteration in network components. The dysregulation or alteration of network components or networks can be achieved either by knocking down one or more nodes or rewiring their connections. Thus, it assesses any significant associations between a network’s structural or topological properties and functional essentiality. Further, network vulnerability analysis is among the most efficient approaches of network analysis, which assists in identifying key molecular players based on various topological properties of a network at a system level. Herein, we present a software tool named NetVA, implemented and developed using the R programming language as an R package for the vulnerability and influence analysis of a protein-protein interactions network (PPIN). The NetVA package is faster in analysis than other similar tools because it utilizes the parallel processing features of the parallel package implemented in base R to speed up the network analysis process. However, this feature of the NetVA could be achieved only on macOS or Linux-based operating systems due to the limitations of the underlying R package.

To demonstrate the applicability and usefulness of the package, a PPIN of breast cancer having 143751 interactions among 10597 proteins was used and analyzed using NetVA. As a result, a list of vulnerable proteins (VPs) was obtained by considering a minimum of three topological properties based on the presence of nodes (proteins) in the list of outliers as considered for the analysis. From the lists of VPs obtained from both approaches, it was observed that non-essential proteins, including non-hubs and non-bottlenecks, were also detected as vulnerable or critical, along with essential, hub, and bottleneck proteins. The commonly identified influential proteins (IPs) based on EVC and EVC+ rankings, coupled with the Pareto principle of the 80:20 rule, were also found to have some experimentally validated essential genes. Furthermore, it was noticed that several VPs and IPs, including APP, BCL2, BRCA1, ERBB2, ESR1, JUN, and TP53, are well-known for their role in the pathogenesis and progression of breast carcinoma. For example, the APP (Amyloid precursor protein) has been reported to promote breast cancer cell migration and invasion by regulating the MAPK signaling pathway.^38^ Next, the BCL2 (B-cell lymphoma 2) regulates cell death (apoptosis), either by inducing or inhibiting the apoptotic process. However, the gene encoding BCL2 could promote cell proliferation due to its downregulation in invasive breast tumors.^39^ Next, the *BRCA1* encoding Breast cancer type 1 susceptibility protein acts as a tumor suppressor and is associated with response to DNA damage stimulus. The *BRCA1* gene has been reported to be a mutated candidate gene for breast cancer.^40^ Moreover, the gene encoding ERBB2 is overexpressed or amplified in approximately 30% of cases of human breast cancer.^41,42^ Further, its overexpression is associated with increased metastasis and resistance of cancer cells to cancer therapies such as chemotherapy, hormone therapy, and radiation therapy. Next, the *ESR1* gene encoding Estrogen receptor alpha is frequently amplified in the case of breast cancer.^43^ Also, the overexpression of *JUN* has been reported to be associated with liver metastasis of human breast cancer.^44^ However, the *TP53* gene acts as a tumor suppressor, regulating uncontrolled cell growth and division. It has also been reported to be a mutated cancer gene in the case of breast cancer.^40^

When vulnerable protein pairs (VPPs) were identified from all the VPs, it resulted in some interesting findings. For example, among all VPPs, some protein pairs were of synthetic lethality in nature when checked with a comprehensive list of known synthetic lethal gene pairs. Synthetic lethality indicates those genetic interactions in which, when two genes are simultaneously perturbed, it leads to the death of a cell or organism.^45-47^ Thus, all the VPPs identified in the present study, especially those showing synthetic lethal interactions, are of key interest, which may help to understand breast cancer biology in a better way and help toward the development of novel and efficient therapies.

The results discussed above suggest that the NetVA package is helpful in identifying potential novel biomarkers or therapeutic candidates. For the vulnerability analysis of networks containing 10,000 or more nodes, a large amount (i.e., several dozen of gigabytes) of memory is required for simulating networks after the deletion of each node one by one and calculating various topological properties to check the effect of deleted or knocked-out nodes, node pairs or node triplets on the resultant networks. The same computational resource is also required for the EVC and EVC+-based analysis of large networks. The present analysis used the R statistical software with the parallel and igraph R packages on a 64-bit Ubuntu computer with a 40-core processor and 95 GB physical memory. However, the network vulnerability and influence analysis can also be carried out on a computational resource with less physical memory and processing capability.

The PPI data produced as a result of experimental techniques are incomplete and have noise. Also, few proteins or pathways which are highly studied are well annotated and hence exhibit in-depth information, including their functions and interactions with other molecules. This may result in significant biases in the construction of PPI networks and, consequently, a reduction in the accuracy of network analysis methods to predict essential proteins or VPs. However, the present study did not aim to reconstruct the investigated networks accurately; the problem of dealing with this bias-ness needs further investigation. Therefore, there is a big challenge to consider for further research on developing and presenting effective pre-processing techniques suitable for each condition or organism while examining the robustness of biological networks to process PPI data for subsequent use.

## Conclusion

The NetVA provides an easy-to-use and time-efficient R-based tool for network vulnerability and EVC-based analysis with the utilization of parallel computing implemented in the R statistical computing environment. It provides an effective, system-level approach for analyzing complex biological networks to decipher the most critical or vulnerable molecules (single proteins, protein pairs, and protein triplets), which can further be utilized to develop more efficient diagnostic and therapeutic strategies for better disease management. Additionally, it can be applied to any networks, including biological such as disease, process, and species-specific protein interactions networks, to identify vulnerable key molecules or network components with the potential to affect the structural and functional behavior of the respective systems or networks.

## Acknowledgments

VV would like to thank the Indian Council of Medical Research (ICMR), New Delhi (ISRM/12(72)/2020, ID: 2020-2951), Institution of Eminence – University of Hyderabad (No. UoH/IoE/RC3-21-052), and Department of Biotechnology (DBT) – Government of India (GoI), New Delhi (No. BUILDER-DBT-BT/INF/22/SP41176/2020) for their financial support. SK also acknowledges the University of Hyderabad for Non-NET Fellowship and ICMR for Senior Research Fellowship (Grant No.: 3/2/2/113/2019/NCD-III, ID: 2019-6723). The authors would like to thank the Center for Modeling, Simulation & Design (CMSD), University of Hyderabad, for providing computational facilities.

## Conflict of interest

The authors declare no conflict of interest.

## Supporting Information

**Table S1**. Topological properties of all nodes, i.e., proteins present in the network.

**Table S2**. Distribution of vulnerable proteins based on a minimum of three properties.

**Table S3**. Common Influential genes based on EVC and EVC+ values.

**Table S4**. List of proteins that are vulnerable as well as influential.

**Table S5**. Topological properties of protein pairs (edges) present in the network.

**Table S6**. Distribution of vulnerable protein pairs based on a minimum of three properties.

## Notes

### Competing Interest Statement

The authors have declared no competing interest.

## References

1. Alberts B. (1998) The cell as a collection of protein machines: preparing the next generation of molecular biologists. Cell, 92: 291–294.

2. Gavin A.-C. et al. (2006) Proteome survey reveals modularity of the yeast cell machinery. Nature, 440: 631–636.

3. Rachita HR, Nagarajaram HA. (2014) Viral proteins that bridge unconnected proteins and components in the human PPI network. Molecular BioSystems, 10(9): 2448–58.

4. Rai A, Pradhan P, Nagraj J, Lohitesh K, Chowdhury R, Jalan S. (2017) Understanding cancer complexome using networks, spectral graph theory and multilayer framework. Scientific reports, 7, 41676. https://doi.org/10.1038/srep41676

5. Kumar S, Lata KS, Sharma P, Bhairappanavar SB, Soni S, Das J. (2019) Inferring pathogen-host interactions between Leptospira interrogans and Homo sapiens using network theory. Scientific reports, 9(1): 1–7.

6. Carter S, Brechbuler C, MGriffin Bond A. (2004) Gene co-expression network topology provides a framework for molecular characterization of cellular state. Bioinformatics, 20 (14): 2242–2250.

7. Srivastava A, Kumar S, Ramaswamy R. (2014) Two-layer modular analysis of gene and protein networks in breast cancer. BMC systems biology, 8 (1):1–6.

8. Li J, Zhou D, Qiu W, Shi Y, Yang JJ, Chen S, et al. (2018) Application of weighted gene co-expression network analysis for data from paired design. Scientific reports, 8 (1): 1–8.

9. Li CY, Cai JH, Tsai JJ, Wang CC. (2020) Identification of hub genes associated with development of head and neck squamous cell carcinoma by integrated bioinformatics analysis. Frontiers in oncology, 10, 681.

10. Dartnell L, Simeonidis E, Hubank M, Tsoka S, Bogle IDL, Papageorgiou LG. (2005) Robustness of the p53 network and biological hackers. FEBS Lett., 579 (14), 3037–3042

11. Friedel CC, Zimmer R. (2007) Influence of degree correlations on network structure and stability in protein–protein interaction networks. BMC Bioinform., 8, 297

12. Abdi A, Tahoori MB, Emamian ES. (2008) Fault diagnosis engineering of digital circuits can identify vulnerable molecules in complex cellular pathways. Sci. Signal, 1 (42), ra10.

13. Zotenko E, Mestre J, O’Leary DP, Przytycka TM. (2008) Why do hubs in the yeast protein interaction network tend to be essential: reexamining the connection between the network topology and essentiality. PLoS Comput. Biol., 4 (8), e1000140.

14. Zheng H, Wang H, Azuaje F. (2010) eNelator: A simulation system for large-scale vulnerability analysis of species-, disease-and process-specific protein networks. Journal of Computational Science, 1(4), 197–205.

15. Ullah, A., Wang, B., Sheng, J. et al. (2022) Escape velocity centrality: escape influence-based key nodes identification in complex networks. Applied Intelligence, 1–19.

16. Jeong H, Mason SP, Barabasi AL, Oltvai ZN. (2001) Lethality and centrality in protein networks. Nature, 411: 41–10.

17. Bartha I, di Iulio J, Venter J., et al. (2018) Human gene essentiality. Nat Rev Genet, 19: 51–62.

18. Meng X, Wang J, Yuan C., et al. (2015) CancerNet: a database for decoding multilevel molecular interactions across diverse cancer types. Oncogenesis 4, e177.

19. Raj, S.; Anil, A.P.; Shukla, A.; Anoosha, K.; Srivastava, A. Benchmark gene reference data for Breast Cancer. Mendeley Data, V2, 2022. doi: 10.17632/xdkvk75ns7.2

20. Raj, S; Anil, AP; Shukla, A; Anoosha, K; Srivastava, A. Benchmark data set for breast cancer associated genes. Data in Brief, 2022, 45, 108583.

21. R Core Team. (2021) R: A language and environment for statistical computing. R Foundation for Statistical Computing, Vienna, Austria, URL https://www.R-project.org/.

22. Ritchie ME, Phipson B, Wu DI, Hu Y, Law CW, Shi W, Smyth GK. (2015) limma powers differential expression analyses for RNA-sequencing and microarray studies. Nucleic acids research, 43 (7): e47–e47.

23. Love MI, Huber W, Anders S. (2014) Moderated estimation of fold change and dispersion for RNA-seq data with DESeq2. Genome biology, 15 (12): 1–21.

24. Robinson MD, McCarthy DJ, Smyth GK. (2010) edgeR: a Bioconductor package for differential expression analysis of digital gene expression data. Bioinformatics, 26 (1):139–40.

25. Charif D, Lobry JR. (2007) SeqinR 1. 0-2: a contributed package to the R project for statistical computing devoted to biological sequences retrieval and analysis. Structural approaches to sequence evolution. Springer, Berlin, Heidelberg, 207–232.

26. Csardi G, Nepusz T. (2006) The igraph software package for complex network research. InterJournal, complex systems, 1695 (5): 1–9.

27. Newman M E J. (2005) Power laws, Pareto distributions and Zipf’s law. Contemp Phys, 46, 323–351.

28. Dong J, Horvath S. (2007) Understanding network concepts in modules, BMC Syst. Biol. 1, 24.

29. Albert R, Jeong H, Barabasi AL. (2000) Error and attack tolerance of complex networks. Nature, 406 (6794): 378–382.

30. Han JD, Bertin N, Hao T, Goldberg D, Berriz G, Zhang L, Dupuy D, Walhout A, Cusick M, Roth F, Vidal M. (2004) Evidence for dynamically organized modularity in the yeast protein-protein interaction network. Nature, 430 (6995): 88–93.

31. Carlson M, Zhang B, Fang Z, Mischel P, Horvath S, Nelson SF. (2006) Gene connectivity, function, and sequence conservation: Predictions from modular yeast coexpression networks. BMC Genomics, 7 (1): 1–15.

32. Horvath S, Zhang B, Carlson M, Lu K, Zhu S, Felciano R, Laurance M, Zhao W, Shu Q, Lee Y, Scheck A, Liau L, Wu H, Geschwind D, Febbo P, Kornblum H, Cloughesy T, Nelson S, Mischel P. (2006) Analysis of oncogenic signaling networks in Glioblastoma identifies ASPM as a novel molecular target. Proc Natl Acad Sci USA, 103 (46): 17402–17407.

33. Liang Y, Yin S, Li F, Peng H. (2018) Role of Articulation Point in Complex Networks, International Conference on Robots & Intelligent System (ICRIS).

34. Freeman LC. (1978) Centrality in social networks: Conceptual clarification. Social Networks, 1, 215–239.

35. Watts DJ, Strogatz SH. (1998) Collective dynamics of ‘small-world’ networks. Nature, 393 (6684): 440–2. 10.1038/30918.

36. Ravasz E, Somera AL, Mongru DA, Oltvai ZN, Barabasi AL. (2002) Hierarchical organization of modularity in metabolic networks. Science, 297 (5586): 1551–5.

37. Ma HW, Zeng AP. (2003) The connectivity structure, giant strong component and centrality of metabolic networks. Bioinformatics, 19 (11): 1423–1430.

38. Wu X, Chen S, Lu C. (2020) Amyloid precursor protein promotes the migration and invasion of breast cancer cells by regulating the MAPK signaling pathway, International Journal of Molecular Medicine, 45 (1), 162–174.

39. Tan P, Bay B, Yip G, et al. (2005) Immunohistochemical detection of Ki67 in breast cancer correlates with transcriptional regulation of genes related to apoptosis and cell death. Mod Pathol, 18, 374–381.

40. Sjoblom T, Jones S, Wood LD, et al. (2006) The consensus coding sequences of human breast and colorectal cancers. Science, 314: 268–74.

41. Slamon DJ, Clark GM, Wong SG, Levin WJ, Ullrich A, McGuire WL. (1987) Human breast cancer: correlation of relapse and survival with amplification of the HER-2/neu oncogene. Science, 235(4785):177–182.

42. Slamon DJ, Godolphin W, Jones LA, Holt JA, Wong SG, Keith DE, Levin WJ, Stuart SG, Udove J, Ullrich A, Press MF. (1989) Studies of the HER-2/neu proto-oncogene in human breast and ovarian cancer. Science, 244(4905): 707–712.

43. Holst F, Stahl PR, Ruiz C, Hellwinkel O, Jehan Z, Wendland M, Lebeau A, Terracciano L, Al-Kuraya K, Jänicke F, Sauter G. (2007) Estrogen receptor alpha (ESR1) gene amplification is frequent in breast cancer. Nature genetics, 39 (5): 655–60.

44. Zhang Y, Pu X, Shi M. et al. (2007) Critical role of c-Jun overexpression in liver metastasis of human breast cancer xenograft model. BMC Cancer, 7, 145.

45. Tang S, Gökbağ B, Fan K, Shao S, Huo Y, Wu X, Cheng L and Li L (2022) Synthetic lethal gene pairs: Experimental approaches and predictive models. Front Genet, 13, 961611.

46. Dobzhansky, T. (1946) Genetics of natural populations; recombination and variability in populations of Drosophila pseudoobscura. Genetics, 31(3), 269–290.

47. Lucchesi, JC. (1968) Synthetic lethality and semi-lethality among functionally related mutants of Drosophila melanfgaster. Genetics, 59(1), 37–44.

